# CRISPR-Cas9 mediated endogenous utrophin upregulation improves Duchenne Muscular Dystrophy

**DOI:** 10.1101/2023.04.18.536394

**Authors:** Simon Guiraud, Sumitava Dastidar, Fetta Mazed, Fatima Amor, Maelle Ralu, Anne de Cian, Isabelle Richard, Giuseppe Ronzitti, Francesco Saverio Tedesco, Mario Amendola

## Abstract

Duchenne muscular dystrophy (DMD) is a lethal neuromuscular disorder caused by loss of dystrophin. Upregulation of utrophin (UTRN), a dystrophin paralogue, is a promising therapeutic avenue. Here, we present a CRISPR-Cas9-mediated strategy to increase utrophin expression by disrupting microRNA (miR) binding sites (BS). Using a Cas9/gRNA ribonucleoprotein (RNP) complex we disrupted several miR BS in DMD myoblasts and selected the Let-7c BS has crucial for UTRN repression. Interestingly, Cas9/gRNA indels were as efficient as the complete removal of Let-7c BS in upregulating UTRN expression, without any major off-targets. In three-dimensional human DMD cultures, Cas9/gRNA-mediated editing resulted in significant utrophin upregulation and functional improvements of calcium dysregulation and muscle contraction. Finally, Let-7c BS disruption in mdx animals by systemic rAAVs mediated delivery of Cas9 and gRNA resulted in utrophin upregulation and amelioration of the muscle histopathological phenotype. These findings provide the foundations for a universal (mutation-independent) gene editing therapeutic strategy for DMD.

**One Sentence Summary:** CRISPR-Cas9 has the potential to upregulate utrophin to treat all DMD patients.

## INTRODUCTION

Duchenne muscular dystrophy (DMD, OMIM 310200) (*1, 2*) is a lethal X-linked neuromuscular disorder affecting 1 in 5000 new-born males (*3*). The disease is caused by mutations in the dystrophin gene leading to loss of dystrophin, a large sub-sarcolemmal protein essential to maintain the biomechanical properties of fiber strength, flexibility and stability in skeletal muscle (*4*). Dystrophin forms part of the dystrophin-associated protein complex (DAPC) and establishes a stable link between the extracellular matrix and the actin cytoskeleton allowing myofibers to cope with repeated cycles of muscle contraction and relaxation (*5*). In the absence of dystrophin, the sarcolemma becomes highly susceptible to contraction-induced injuries causing muscle degeneration and replacement of contractile material with adipose and fibrotic tissues. DMD patients manifest the first onset of symptoms in early childhood, lose ambulation generally by the age of 12 (*6*) and develop respiratory and cardiac failure leading to premature death by their early thirties (*7*).

Despite clinical management of cardiac complications, assisted ventilation and corticosteroid treatment (*8, 9*), there is currently no cure for DMD. The urgency to seek a therapy for DMD has resulted in parallel efforts to develop gene-based, cell-based and pharmacological strategies. Among gene-based approaches, exon-skipping, such as Eteplirsen (*10*), Golodirsen (*11*), Viltolarsen (*12*), Casimersen (*13*), and stop codon read-through (Translarna/Ataluren) (*14*) strategies offer limited clinical benefits and are mutation-specific (i.e., only suitable for specific subsets of DMD patients). Gene therapy using recombinant associated adenovirus (rAAV) delivering a truncated and partially functional micro-dystrophin recently entered in clinical trials, bringing hope of an effective therapy for DMD (*15*). An alternative therapeutic approach, potentially suitable to all DMD patients irrespective of their genetic defect, consists in upregulating utrophin, a structural and functional paralogue of dystrophin, able to compensate for the dystrophin deficit (*16–18*). The utrophin gene exhibits 65% nucleic-acid identity and an intron-exon structure very similar to the dystrophin gene (*19*), suggesting that the two genes arose through an ancient duplication event. The encoded 394 kDa protein contains a modular organization similar to dystrophin with an actin-binding N-terminus, a triple coiled-coil spectrin repeat central region, and a C-terminus that consists of protein-protein interaction motifs and shares many binding partners, such as α-dystrobrevin-1, β-dystroglycan and F-actin (*4, 20*). However, utrophin differs from dystrophin in its mode of interaction with microtubules (*21*) and actin filaments (*22*) and cannot anchor nNOS (*23*). The spatio-temporal expression is also different, as utrophin is ubiquitously localized at the sarcolemma in utero, progressively replaced by dystrophin (*24, 25*) and enriched in adult muscles at the neuromuscular and myotendinous junctions (*26*) as well as the sarcolemma of regenerated myofibers (*27*).

Despite these differences, utrophin can act as a surrogate to compensate for the lack of dystrophin in DMD (*28*). Seminal studies with transgenic mice overexpressing full-length utrophin in skeletal muscles established that its expression suppresses functional signs of dystrophy in a dose-dependent manner (*29*) without any toxicity (*30*) in the mdx mice, a widely used animal model of DMD (*31*). While a 1.5-fold increase of utrophin results in a therapeutic benefit, a 3-4-fold overexpression prevents the dystrophic pathology (*29, 32*). Moreover, vector delivery of a truncated utrophin minigene partially prevents the pathology in dystrophic mice (*33, 34*) and GRMD dogs (*35*), with a favorable immunologic profile compared to truncated dystrophin (*36, 37*). Overall, these studies in animal models of DMD strongly support utrophin as a functional surrogate for dystrophin and emphasize the potential of utrophin upregulation for the treatment DMD. Therefore, several strategies to modulate utrophin have been proposed (*38*). Small drugs increase utrophin expression at transcriptional level and results in functional improvements in animal models (*39, 40*). Unfortunately, Ezutromid/SMT C1100, a first-in-class orally bioavailable small utrophin modulator, showed clinical benefit in interim results in Phase 2 trials in DMD patients (NCT02858362), but failed to meet its primary endpoints due to a lack of sustained efficacy attributed in part to hepatic metabolism and clearance (*39, 41*). In addition to transcriptional control, utrophin expression is post-transcriptionally regulated by several miRs, such as miR-150, -296b, -133b or Let-7c, able to interact with its 3’ untranslated region (UTR) (Fig. 1) (*42*). Masking miR Let-7c BS with 2′-O-methyl-phosphorothioate (2OMePS) (*43*) and peptide conjugated to phosphorodiamidate morpholino oligomers (P-PMO) (*44*) blocks miR-mediated repression and results in transient utrophin upregulation. Although these strategies represent a novel promising therapeutic avenue, they would require life-long repeated administration.

**Fig. 1.**
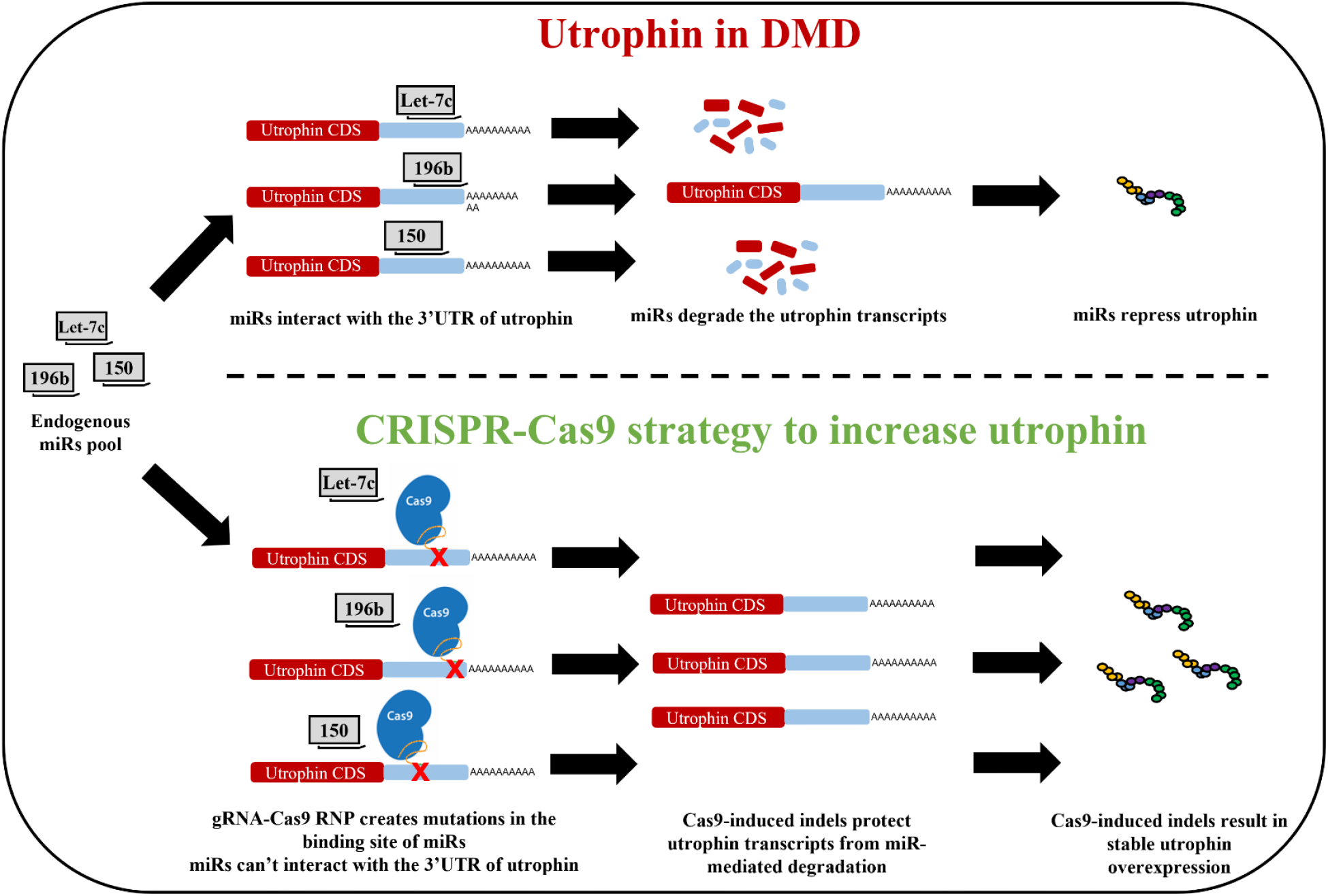
Schematic representation of the CRISPR-Cas9 strategy to disrupt miR BS for utrophin upregulation. miRs such as Let-7c, 196b or 150 have the potential to bind the 3’UTR of utrophin mRNA, degrade transcripts and repress translation. CRISPR-Cas9 can be used to create mutation in miR BS, thus preventing their interaction with miRs and degradation of utrophin transcripts, resulting in utrophin upregulation.

In recent years, the clustered regularly interspaced short palindromic repeats (CRISPR) and CRISPR-associated (Cas) protein system has emerged as a revolutionary gene editing tool to permanently and precisely edit any DNA locus and correct genetic errors in a wide range of diseases (*45, 46*). Several groups demonstrated that the CRISPR-Cas9 system was able to correct and restore dystrophin expression in immortalized DMD patient cells (*47*) as well as in mdx (*48–50*) and GRMD (*51*) animal models of DMD. These approaches are mutation specific and could induce immune response to the newly expressed dystrophin (*52*). Alternatively, utrophin upregulation was successfully achieved by constitutive expression of a ‘dead’ Cas9 (dCas9) nuclease fused to transcription enhancing factors able to activate the utrophin promoter without cleaving DNA (*53, 54*). Although promising, this system requires stable expression of the activator. Very recently, CRISPR-Cas9 was used in human (*55*) and murine (*56*) cells to remove several miR BS from the 3’UTR of the endogenous utrophin gene. This hit-and-run approach resulted in a 1.5-2-fold *in vitro* utrophin upregulation: however, no *in vivo* or functional data were provided.

In the present study, using the CRISPR-Cas9 system, we systematically disrupted the BS of such miRs at the DNA level to permanently upregulate utrophin (Fig. 1). We first established that Let-7c is the main miR responsible of utrophin post-transcriptional downregulation and that a minimal disruption of its BS was sufficient to upregulate utrophin in murine and human myoblasts and myotubes. Noteworthy, BS disruption also resulted in clear functional benefits in three-dimensional (3D) engineered human DMD muscles, Finally, we demonstrated utrophin upregulation and amelioration of the muscle histopathological phenotype in mdx mice. These data demonstrate the *in vitro* and *in vivo* potential of the proposed CRISPR-Cas9 strategy to upregulate utrophin expression as a mutation-independent therapeutic strategy for all DMD patients.

## RESULTS

### Evaluation of gRNAs targeting miR BS for utrophin gene upregulation in hDMD myoblasts

To identify genomic sequences involved in post-transcriptional repression of utrophin, we designed gRNAs targeting previously described miR BS present in the 3’UTR of utrophin, i.e. the miR-150/133b/296, miR-196b and miR-Let-7c 5p (Table. S1) (*42*). SpCas9/gRNA ribonucleoprotein (RNP) complexes were transfected in skeletal myoblasts derived from a DMD patient with a deletion of the dystrophin exon 52 (Δ52). After 7 days, we quantified the Cas9 indels rate and pattern of each gRNA (Fig 2 A-C and Fig S1). The gRNA targeting miR-150/133b/296 BS demonstrated low indels (14% ±1.3 SEM, which could explain the lack of increase of utrophin mRNA (Fig. 2D). Despite high cutting efficacy (63% ±3.5 SEM), the gRNA targeting the miR-196b only minimally increased utrophin mRNA suggesting that this BS is not a good candidate to pursue (Fig. 2D). Finally, all gRNAs targeting Let-7c resulted in >50% indel efficacy (Fig. 2A) and significantly upregulated utrophin mRNA level (Fig. 2D), highlighting the importance of Let-7c BS in controlling utrophin expression. Surprisingly, gRNA Let-7c-1 resulted in consistently higher utrophin mRNA upregulation compared to other gRNAs and its effect was confirmed at protein level (Fig. 2E). This difference could be due to the different indel patterns created by each gRNA. While the indels generated by the gRNA hLet-7c-2 do not affect miRNA seed (Fig S1E), gRNAs hLet-7c-1 and hLet-7c-3 are close or delete part of the Let7-c BS seed sequence, which is crucial for miR BS (Fig. 2C) (*57*). Although they were shifted of only 3 nucleotides and gave similar indels, mostly generating a +1 nucleotide insertion and -8 nucleotides deletion (Fig. 2B; Fig S1E), gRNA Let-7c-1 was more efficient than gRNA Let-7c-3 in upregulating utrophin, suggesting that even small differences can have profound effect on miR BS.

**Fig. 2.**
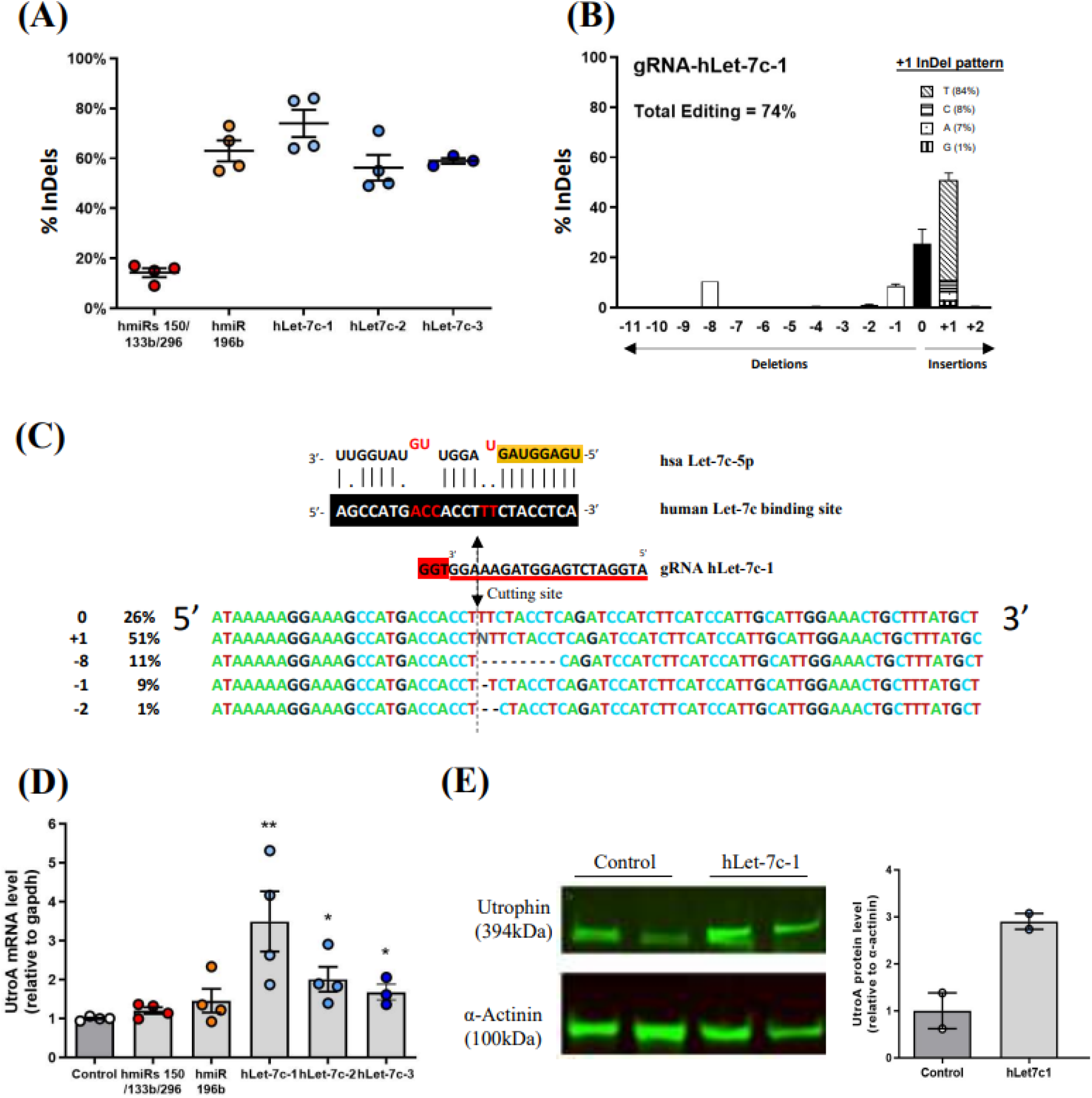
Screening of gRNAs to upregulate utrophin expression in hDMD Δ52 myoblasts. hDMD Δ52 myoblasts were transfected with different Cas9/gRNA RNP and gDNA and protein were collected at days 7 for analyses. **(A)** TIDE quantification of the indels generated by the gRNAs targeting the indicated miRs-150/133b/296, 196b and Let-7c BS on the 3’UTR of utrophin. Bars are mean ± SEM of n = 4 per condition. Dots represent each experiment. **(B)** ICE indel profiles generated by the gRNA-hLet-7c-1. In the inset are indicated the % at which each nucleotide is inserted in the +1 indel. Bars are mean ± SEM of n = 4 per condition. **(C)** Top: representation of the interaction of the miR hLet-7c-5p with its BS on the 3’UTR of utrophin. Seed region of miR Let-7c-5p is highlighted in yellow, while mismatching nucleotides are indicated in red. Localization and cutting site of the gRNA hLet-7c-1 is specified (arrow). The NGG PAM site is highlighted in red. Bottom: representative InDel sequence with ICE software. On the left is indicated the frequency of each indel. **(D)** Quantitative PCR analysis of utrophin A mRNA expression level, normalized with gapdh, after editing with the indicated gRNAs. Bars are mean ± SEM of n = 4 per condition. Dots represent each experiment. *p < 0.05, **p < 0.01. **(E)** Relative utrophin protein expression after editing with the indicated gRNA was determined by western blot and standardized for α-actinin loading. Bar graph: WB quantification. Relative utrophin expression is shown as mean ± SEM of n = 2 per condition. Dots represent each experiment.

Overall, these data demonstrate that CRISPR-Cas9-mediated Let-7c BS disruption results in ∼3 fold utrophin upregulation in DMD myoblasts.

### Scanning utrophin 3’UTR for post-transcriptional regulatory sequences

Next, we sought to identify additional sequences present in human utrophin 3’UTR that could affect mRNA stability and translation. Based on previous data generated on mouse utrophin 3’UTR (*58, 59*), we predicted several regulatory sequences (Fig 3 A) and we generated seven 3’UTR variants with deletions encompassing one or more of these sequences. Each variant, as well as the wild type sequence as reference, was inserted at the 3’ UTR of the Gaussia luciferase (GLuc) reporter gene in dual reporter plasmid containing the secreted embryonic alkaline phosphatase (SeAP) as normalizer for transfection efficacy (Fig 3 A-B). These plasmids were transfected in hDMD Δ52 myoblasts and GLuc and SeAP levels were quantified 48 hours post-transfection.

**Fig. 3.**
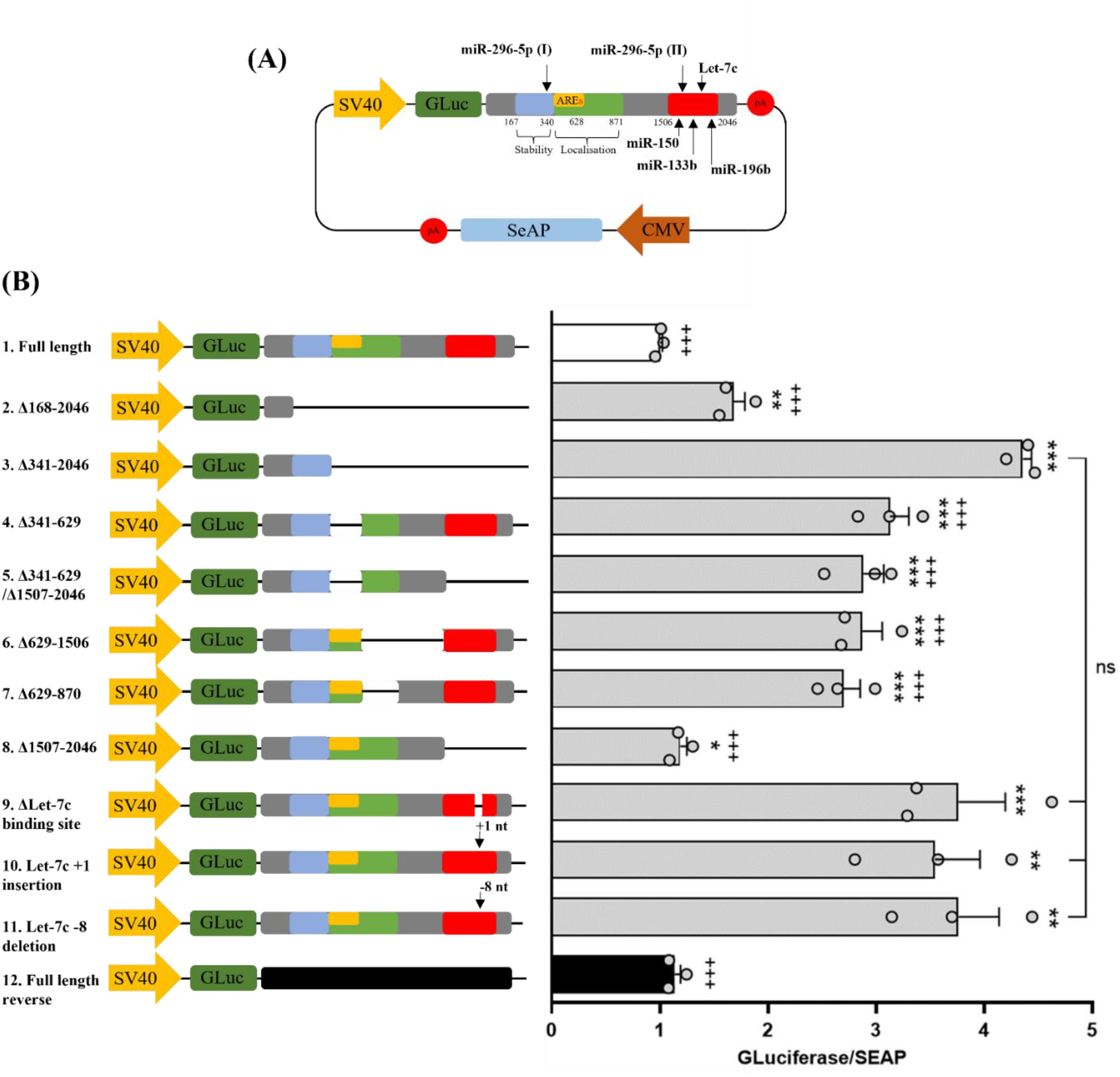
Identification of post-transcriptional regulatory sequences within utrophin 3’UTR. **(A)** Scheme of a dual reporter plasmids. To study the impact of several modifications of the 3’UTR of utrophin, several 3’UTR variants were integrated after the Gaussia luciferase (Gluc) gene, driven by an SV40 promoter. The secreted alkaline phosphatase (SEAP) expression, under the control of a CMV promoter, was used for transfection efficacy and normalization. miR BS and stability, localisation and AU-Rich element predictions are indicated with their nucleotide positions. **(B)** The reporter constructs were transfected in hDMD myoblasts and Gluc and SeAP expression measured 48hrs post-transfection. Left; 3’UTR variants used in this study. For each variant it is indicated the nucleotide position of the deletion. Right: ratio of Gluc/SEAP was calculated for each contract. The reporter construct 1 (Full length) was used to determine the basal level of Gluc expression and construct 12 (Full length reverse) as negative control. Bars represent mean ± SEM of n = 3 per condition. Dots represent each experiment. *p < 0.05, **p < 0.01, ***p < 0.001 versus full length; +p < 0.05, ++p < 0.01, +++p < 0.001 versus D341-2046 3’UTR variant.

Although to different extent, all tested variants resulted in a significant increase of GLuc expression compared to wild type 3’UTR sequence. Either variant 2 or 3 were depleted of the AU-rich elements (AREs), the miR cluster and potential other repressor BS; however, GLuc expression for variant 3 was higher, indicating that the 168-341 sequence is essential to promote stability of utrophin mRNA (*58*). Comparison of variant 4 vs full length confirmed the role of predicted AREs in decreasing utrophin mRNA stability (*60, 61*). Surprisingly, variant 5, which corresponds to variant 4 without the miR BS cluster, did not show any improvement. This was further confirmed with variant 8, where we removed only the miR BS cluster from the full length and we observed only minimal GLuc increase (*56*). A possible explanation could be that the terminal part of the 3’UTR contains both repressing miR BS as well as mRNA stabilizing elements.

To further evaluate the effect of miR Let-7c BS on utrophin expression, we generated the additional variant 9, where only this BS was removed. Intriguingly, this small deletion upregulated GLuc expression (3.8-fold± 0.3 SEM) only slightly less than the best variant 3 (4.4-fold± 0.1 SEM), which contains a ∼1700 bp deletion. Once more, this finding indicates the crucial role of miR Let-7c in regulating utrophin expression. To evaluate if the indels generated by gRNA Let-7c-1 would completely abolish Let-7c binding, we generated two additional 3’UTR constructs each incorporating one of the 2 main indels generated by gRNA hLet-7c-1 (Fig 2 C): a +1 nucleotide (T; variant 10) and a -8 nucleotides deletion (variant 11) (Fig. 3B). Interestingly, with these two 3’UTR variants, we obtained a gene reporter expression similar to ΔLet-7c, variant 9, suggesting that these mutations are both sufficient to completely disrupt Let-7c BS and restore gene expression.

Although we selected our gRNA Let7c-1 for having high specificity (≥3 mismatching nucleotides to other genomic loci) before exploring further this gRNA, we analyzed its potential off-targets. We performed unbiased genome wide Guide-seq analysis, which is known to have low false-positive rate (*62*), in both 293T and DMD Δ52 myoblast human cell lines (Fig. S. 4A-B). Although ranked differently, most of the detected off-targets were found in both cell lines, confirming the importance of the gRNA sequence over the nature of the cell line used. Importantly, only few off-targets had a read abundance >10% of the on-target and all, except one, were targeting intronic or intergenic sequences, limiting the risk of a functional alteration. Top 10 identified off-targets were further analyzed by amplicon-seq in edited hDMD Δ52 myoblasts (Fig. S4C).

In summary, these data demonstrate that our CRISPR-Cas9 strategy targets utrophin 3’UTR with high efficiency and without major off-target effect. Additional studies to reduce off-targets (*63*) and indel quantification on edited muscles, normally exposed to lower gRNA/Cas9 concentration, will be performed before clinical translation.

### Disruption of Let-7c BS results in utrophin upregulation in human DMD myotubes and 3D engineered skeletal muscles

We then evaluated if disruption of Let-7c BS and utrophin upregulation would interfere with the skeletal myogenic differentiation program. 10 days after treatment, we differentiated Cas9 only or Cas9/gRNA treated myotubes (Fig. S2A-B). As expected, mRNA of myogenin, a marker of the late myotubes formation, was increased in myotubes compared to myoblasts (Fig. S2C), while no difference was observed between edited and control myotubes (Fig S2C). We also observed a 2-fold increase of utrophin mRNA in myotubes compared to myoblasts as previously reported (*64*). Importantly, edited myoblasts and myotubes showed a ∼1.8-fold utrophin mRNA upregulation compared to control counterpart (Fig. S2F), indicating that editing mediated utrophin upregulation remained stable during muscle differentiation. Finally, edited myoblast and myotubes showed the same indels percentage and pattern (Fig. S2E-F), suggesting no genotoxicity. Overall, this data demonstrates that this utrophin upregulation strategy is compatible with the physiological myogenic differentiation program.

We then assessed if the editing strategy would induce utrophin upregulation and correct sarcolemma localization in a model more closely resembling skeletal muscle tissue architecture than conventional monolayer cultures (*65*). To this aim, we generated 3D engineered skeletal muscles by culturing Cas9 only or Cas9/gRNA treated hDMD myoblasts in our established 3D platform capable to model muscular dystrophies and therapeutic interventions, including the possibility to study muscle function (*65–69*). 8 days after treatment, edited and control hDMD myoblasts were differentiated into a 3D hydrogel as previously described (*69*) (Fig 4A; Fig S3A-B). After 22-24 days, control and edited 3D muscles showed similar expression of Titin and MyHC as well as similar fusion (percentage of nuclei in fused myotubes out of the total nuclei) and differentiation (percentage of nuclei in MyHC-positive cells) indexes (Fig. S3C-E), indicating similar differentiation potential. Importantly editing remained stable (Fig. 4B-C) and utrophin expression remained upregulated both at mRNA and protein level (Fig. 4D-E). Immunofluorescence staining of cross-sections of the 3D muscles showed once more that utrophin protein was upregulated and correctly localized to the sarcolemma in up to 70% of the edited myofibers compared to control muscle (Figures 4 F-G), providing histological evidence of utrophin protein production in human myofibers in vitro upon CRISPR-Cas9 mediated Let7c BS disruption.

**Fig. 4.**
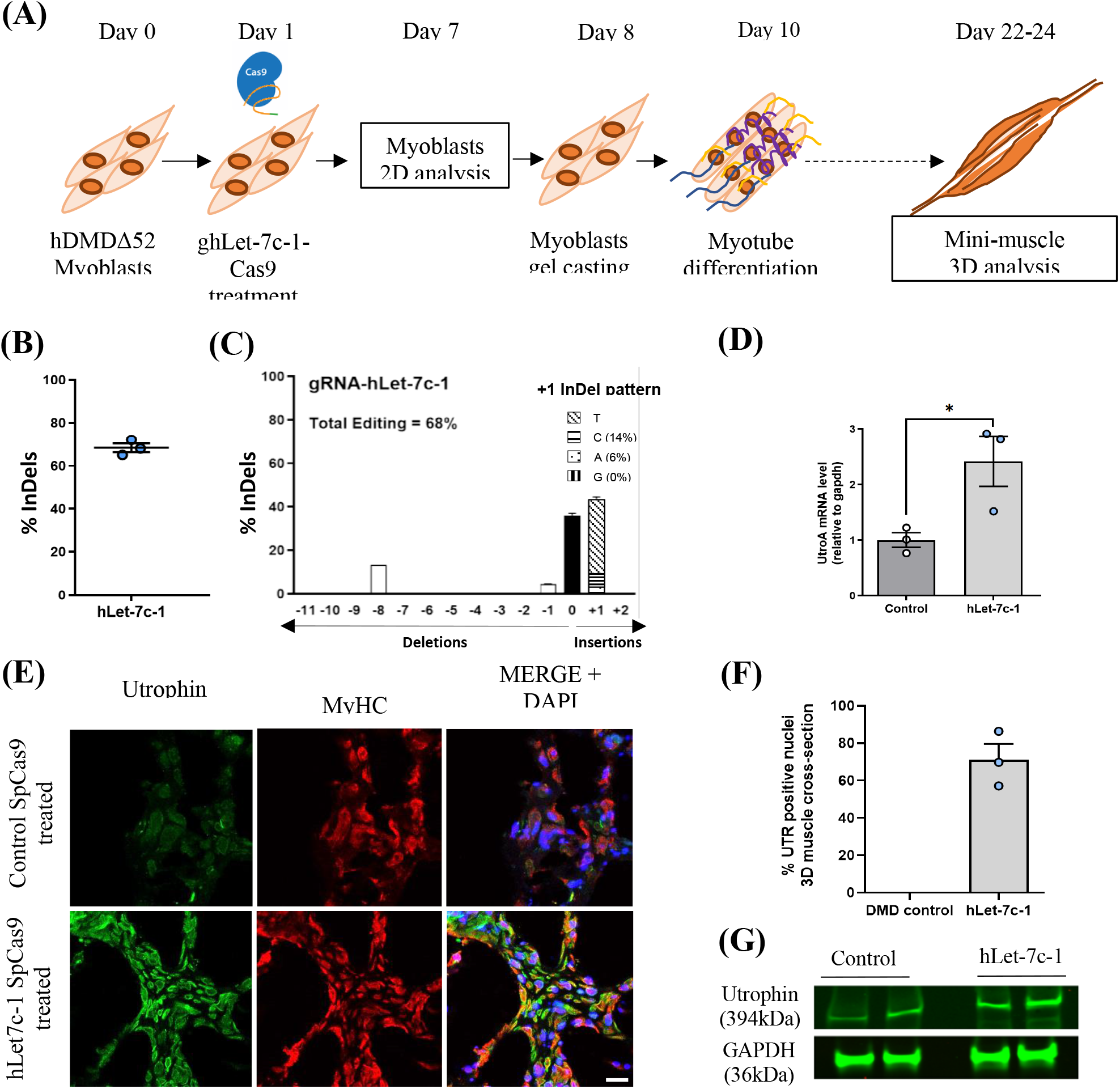
gRNA hLet-7c-1 treatment upregulates utrophin expression in human DMD 3D artificial muscles. **(A)** Schematic representation of the timeline for generating 3D mini muscle starting from treated or untreated myoblasts. **(B)** TIDE quantification of the indels generated by the gRNA Let-7c-1 in 3D hDMD mini-muscle. Bars represent mean ± SEM of n = 3 per condition. Dots represent each experiment. **(C)** Indel profiles generated by the gRNA-hLet-7c-1 in 3D hDMD mini-muscle. In the inset are indicated the % at which each nucleotide is inserted in the +1 indels. Bars are mean ± SEM of n = 3 per condition. **(D)** Quantitative PCR analysis of utrophin A mRNA expression level, normalized with gapdh, after editing with the indicated gRNAs in 3D hDMD mini-muscle. Bars are mean ± SEM of n = 3 per condition. Dots represent each experiment. *p < 0.05. **(E)** Relative utrophin protein expression in 3D DMD muscles after editing with the indicated gRNA was determined by western blot and standardized for GAPDH loading. Bar graph: WB quantification. Bars represent mean ± SEM of n = 2 per condition. Dots represent each experiment. **(F)** Immunofluorescence staining for utrophin and myosin heavy chain (MyHC) on transverse sections of 3D DMD muscles show a higher utrophin signal after treatment with gRNA-hLet-7c compared to control. n = 4 per group. Scale bar: 100 µm. **(G)** Quantification of the immunostaining shown in **(F)**. Bars represent mean ± SEM of n = 3 per condition. Dots represent each experiment.

Overall, these results show that our CRISPR-Cas9 strategy upregulates utrophin expression in human DMD myoblasts and myotubes as well as in 3D engineered DMD skeletal muscles, without affecting myogenic differentiation.

### Utrophin upregulation results in functional benefits in human DMD engineered muscle

In DMD, loss of dystrophin enhances sarcolemma susceptibility to contraction-induced damage. Sarcolemmal lesions and possibly leaky Ca2+ channels increase calcium influx into dystrophic fibers resulting in severe secondary defects such as protease activation, ischemia, mitochondrial dysfunction and metabolic perturbation, ultimately leading to loss of muscle function (*70*). We therefore hypothesized that gRNA hLet-7c-1 mediated utrophin upregulation would result into a measurable amelioration of ‘excitation/contraction coupling’ functional parameters in engineered muscles. To this aim, we performed a calcium transient assay and a whole muscle contractility assay. Upon 10V electrical stimulation, we observed a significant 64% (±14.4 SEM) reduction in intracellular calcium concentration within the hDMD mini-muscles compared to wild-type (wt), demonstrating a clear calcium perturbation (Fig. 5A-B). Following treatment, the intracellular calcium concentration showed a 45% (± 8.2 SEM) calcium increase compared to untreated DMD muscles (Fig. 5A-B). Finally, we assessed the benefit of utrophin upregulation on muscle contraction activity. Unlike wt mini-muscles, efficient muscle contraction was observed in only 57% of the DMD muscles compared to 78% of the edited ones (Fig. 5C).

**Fig. 5.**
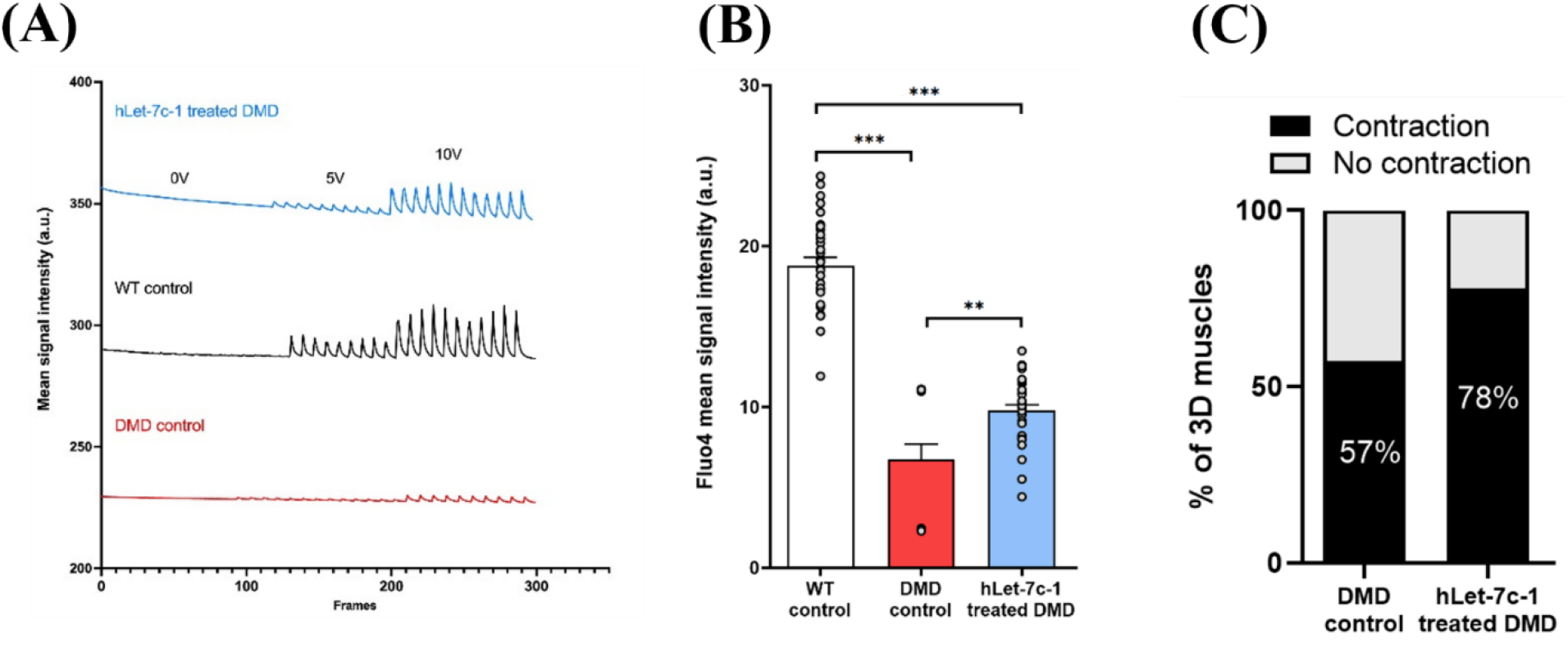
Assessment of calcium dynamics and contractility in gRNA hLet-7c-1 treated DMD 3D muscles. **(A)** Representative results of a calcium transient assay performed on 3D healthy, DMD and gRNA hLet-7c-1 treated DMD mini-muscle. 3D muscles were incubated with media containing Fluo-4AM for 30mins and then washed for 10mins. After 10 seconds without stimulation, 3D muscles were then stimulated using carbon electrodes at 5V for 30sec followed by a 10V stimulation. Fluorescence excitation and emission were collected using a Zeiss 880 confocal LSM to measure signal intensity. Unlike WT control, 3D muscle demonstrate a low signal intensity highlighting calcium influx/efflux perturbations. After treatment with Cas9/gRNA, the calcium transient is rescued toward the WT profile. **(B)** Following a 10V stimulation, hDMD 3D muscle show a significant reduction in Fluo4 mean signal intensity highlighting calcium defects. After editing, the mean signal intensity is significantly improved toward WT levels. Bars represent mean ± SEM. Each dot represents the amplitude of a peak signal obtained upon stimulation. ** p < 0.01; ***p < 0.001. **(C)** Bar graph depicting the percentage of untreated and gRNA hLet-7c-1 treated DMD artificial muscle capable to contract upon electrical stimulation.

Taken together, these results highlight that gRNA hLet-7c-1 mediated utrophin upregulation significantly improved calcium intake and muscle function.

### Evaluation of gRNAs targeting Let-7c BS for utrophin gene upregulation in murine myoblasts

To further evaluate our gene therapy strategy in vivo, we decided to move to the mdx mouse model of DMD (*31*). We first checked for interspecies evolutionary sequence conservation. The miR Let-7c-5p is fully conserved between human and mouse and the miR Let-7c BSs have only a single nucleotide of difference (*71*), despite only 78% of sequence homology between the human and murine utrophin 3’UTR sequences.

Therefore, we designed two novel gRNAs targeting murine Let-7c BS, gRNA mLet-7c-2 and mLet-7c-4 (Table. S1), which are homologous to the human gRNA Let-7c-2 and c-3 except for 1 and 2 nucleotide changes, respectively. These gRNAs were tested in the C2C12 murine myoblast cell line and demonstrated efficient indel generation, with patterns that were similar to human gRNA hLet-7c-1, 2 and 3, with mostly +1 insertions and -8 deletions (Fig. 6A-E). Both gRNAs induced utrophin upregulation by ∼2 and 2.5-fold at mRNA and protein levels, for gRNA mLet-7c-2 and gmLet-7c-4 respectively (Fig. 6F-G), confirming that utrophin post-transcriptional regulation is maintained in murine cells. Overall, gRNA mLet-7c-4 led to higher efficiency than gmLet-7c-2, possibly because gmLet-7c-4 has higher indel efficiency and disrupts the seed of miR Let-7c BS (Fig. 6E).

**Fig. 6.**
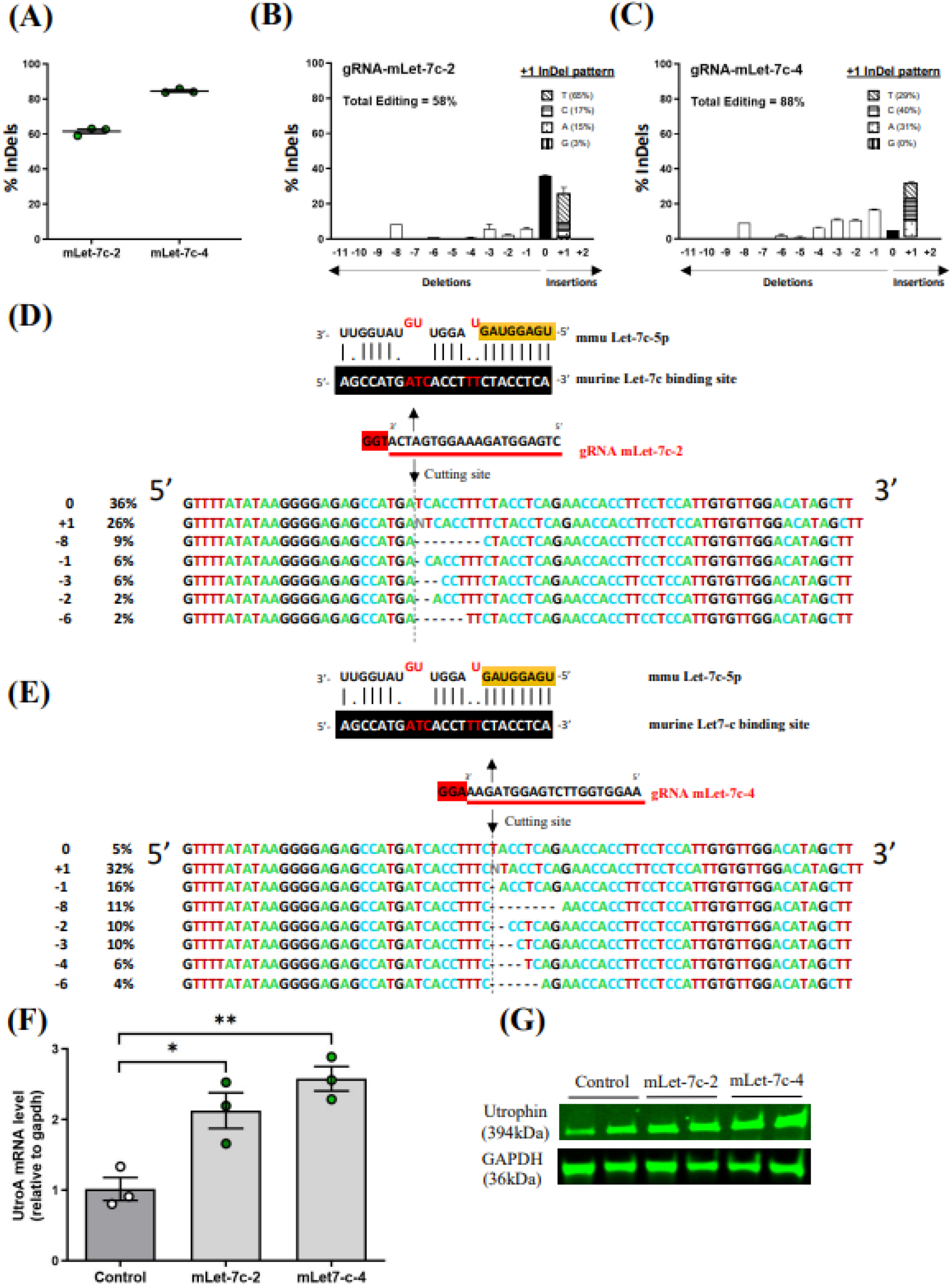
Screening of gRNAs to upregulate utrophin expression in murine C2C12 myoblasts. Murine C2C12 myoblasts were transfected with different Cas9/gRNA and DNA and protein were collected at day 7 for analyses. **(A)** TIDE quantification of the indels generated by the gRNAs targeting mLet-7c BS on the 3’UTR of utrophin. Bars are mean ± SEM of n = 3 per condition. Dots represent each experiment. **(B-C)** Indel profiles generated by the gRNA-mLet-7c-2 and -4. In the insets are indicated the % at which each nucleotide is inserted in the +1 indel. Bars are mean ± SEM of n = 3 per condition. **(D-E)** Top: representation of the interaction of the mviR Let-7c-5 with its BS on the 3’UTR of murine utrophin. Seed region of miR Let-7c-5p is highlighted in yellow, while mismatching nucleotides are indicated in red. Localization and cutting site of the gRNA mLet-7c-2 and -4 are specified (arrows). Their respective NGG PAM sites are highlighted in red. Bottom: representative indel sequence with ICE software. On the left is indicated the frequency of each indel. **(F)** Quantitative PCR analysis of murine utrophin A mRNA expression level, normalized with gapdh, after editing with the indicated gRNAs. Bars are mean ± SEM of n = 4 per condition. Dots represent each experiment. *p < 0.05, **p < 0.01. **(G)** Relative utrophin protein expression after editing with the indicated gRNA was determined by western blot and standardized for GAPDH loading.

In summary, we defined and validated two efficient gRNAs to disrupt Let-7c BS in murine myoblasts and selected gRNA mLet-7c-4 for subsequent in vivo analyses.

### Disruption of Let-7c BS and utrophin upregulation in vivo in mdx mice

To test our strategy in vivo, we generated two rAAV encoding for SpCas9, under the control of the ubiquitous CMV promoter, or gRNA, under the control of the human U6 promoter respectively. We chose the AAV9 serotype, which is known to efficiently target cardiac and skeletal muscles (*72*). To mimic preclinical settings, we systemically injected a rAAV9-CMV-SpCas9 (1.7 10^12^vg) and a rAAV9-U6-gmLet-7c-4 (8.4 10^12^vg) in 4-week-old mdx mice (Fig. 7A), based on previously published work (*73*). As control, we used an rAAV9-U6-gRosa26 encoding for a gRNA targeting the unrelated Rosa26 genomic locus (*49*).

**Fig. 7.**
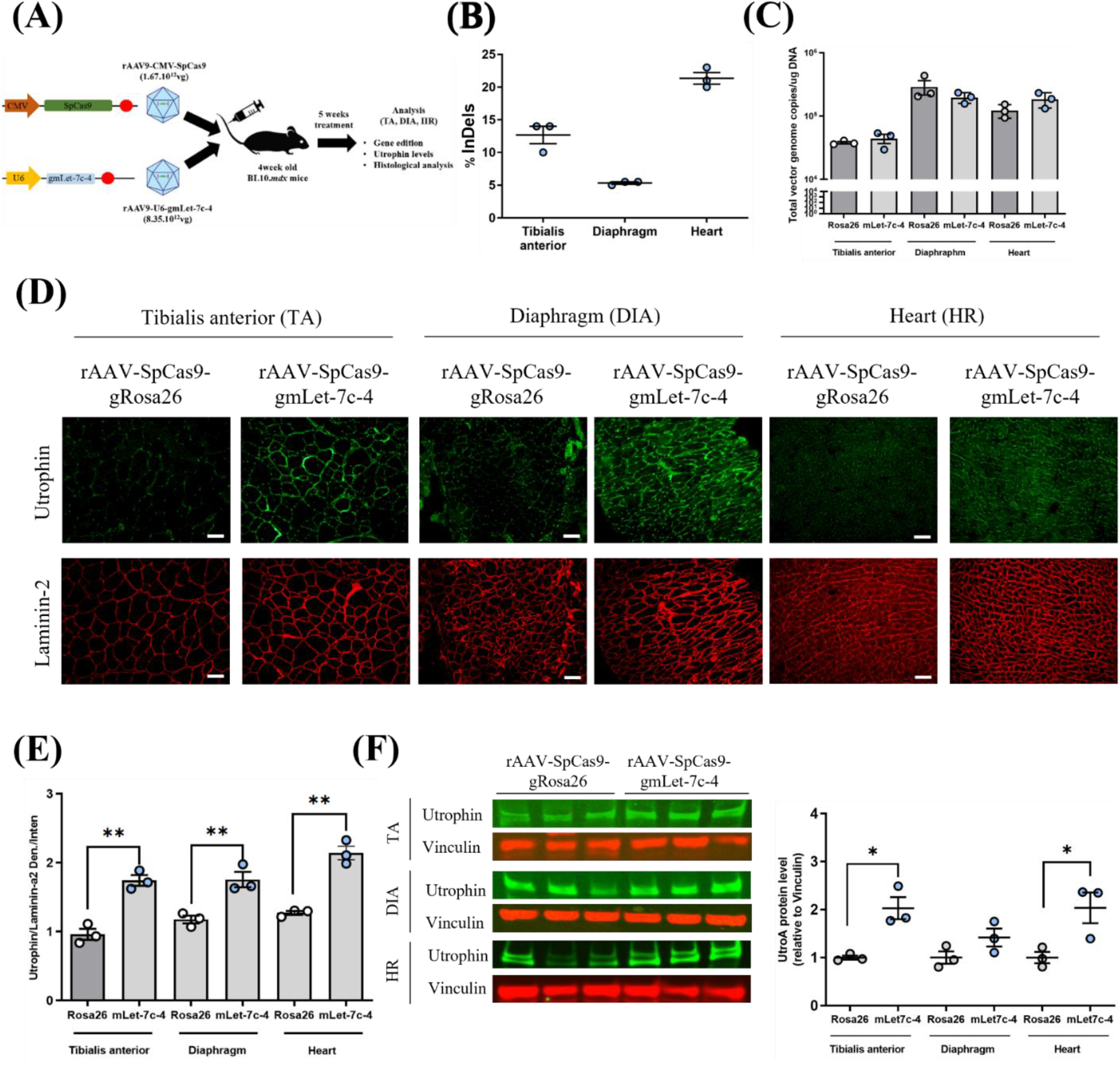
Cas9/gRNA AAV treatment results in upregulation of the endogenous utrophin in *mdx* mice. **(A)** rAAV9-CMV-SpCas9 and rAAV9-U6-mLet-7c-4 were intravenously injected into 4-week-old *mdx* animals. 5 weeks after injection, animals were scarified and downstream analyses performed in tibialis anterior (TA), diaphragm (DIA) and heart (HR) muscles. **(B)** Quantification of rAAV9-CMV-SpCas9 genome copy number in skeletal, respiratory and cardiac muscles of *mdx* mice administered with the indicated AAV encoding gRNAs. Bars are mean ± SEM of n = 3 per condition. Dots represent each mouse. **(C**) NGS quantification of the indels generated by the gRNA mLet7c-4 in TA, DIA and HR muscles after 5 weeks of treatment. Bars are mean ± SEM of n = 3 per condition. Dots represent each mouse. **(D)** Representative images of immunofluorescence of utrophin and laminin-α2 of rAAV-gRNA mLet-7c-4 and rAAV-gRNA Rosa26 (control) treated TA, DIA and HR. Scale bar, 100 μm. **(E)** Quantification of utrophin staining relative to control laminin-α2 of (D). **P < 0.01. **(F)** Relative utrophin and vinculin protein levels from rAAV-gRNA mLet-7c-4 and rAAV-gRNA Rosa26 (control) treated TA, DIA and HR. Bars are mean ± SEM of n = 3 per condition. Dots represent each mouse. TA, tibialis anterior; DIA, diaphragm; HR, heart. *p < 0.05.

After 5 weeks of treatment, mice were sacrificed for analyses. First, we measured rAAV9-CMV-SpCas9transduction efficiency in different muscles. Vector copy numbers were higher in diaphragm (DIA) compared to cardiac (HR) muscles and tibialis anterior (TA) (Fig.7B). We then evaluated on-target editing efficiency for gRNA mLet-7c-4 by next generation sequencing. We achieved ∼13%, 5% and 21% editing in TA, DIA and HR (Figs. 7C, S5). By immunofluorescence, we observed that utrophin signal was consistently increased at the sarcolemma of TA (1.8x ±0.1 SEM), DA (1.5x ±0.1 SEM) and HR (1.7x±0.1 SEM) (Fig. 7D-E). Accordingly, western blot analyses revealed a 2 (±0.2 SEM), a 1.4 (±0.1 SEM) and 2 (± 0.3 SEM)-fold utrophin upregulation in TA, DIA and HR muscles respectively (Fig 7F). Overall, these data demonstrate that our CRISPR-Cas9 strategy upregulates endogenous utrophin levels in both skeletal and cardiac muscles.

To evaluate if this upregulation has an impact on the pathology, we performed histological evaluation of treated mdx mice. In particular, centronucleation, a marker of muscle regeneration (*74*), was significantly reduced by ∼21% (± 1.9 SEM) in TA and by 22% (± 4.0 SEM) in DIA muscle (Fig. 8A, B) in gRNA mLet7c-4 treated mice compared to gRNA Rosa26 treated ones. Areas of necrosis were also significantly reduced by ∼40% (± 0.2 SEM) and 44% (± 0.2 SEM) in skeletal and respiratory muscles after treatment (Fig. 8A, C). Finally, a 10% reduction in fibrosis was observed in the DIA of treated mice (Fig. 8D-E). Amelioration of these histological parameters indicates myofiber protection secondary to utrophin upregulation generated by CRISPR-Cas9 treatment.

**Fig. 8.**
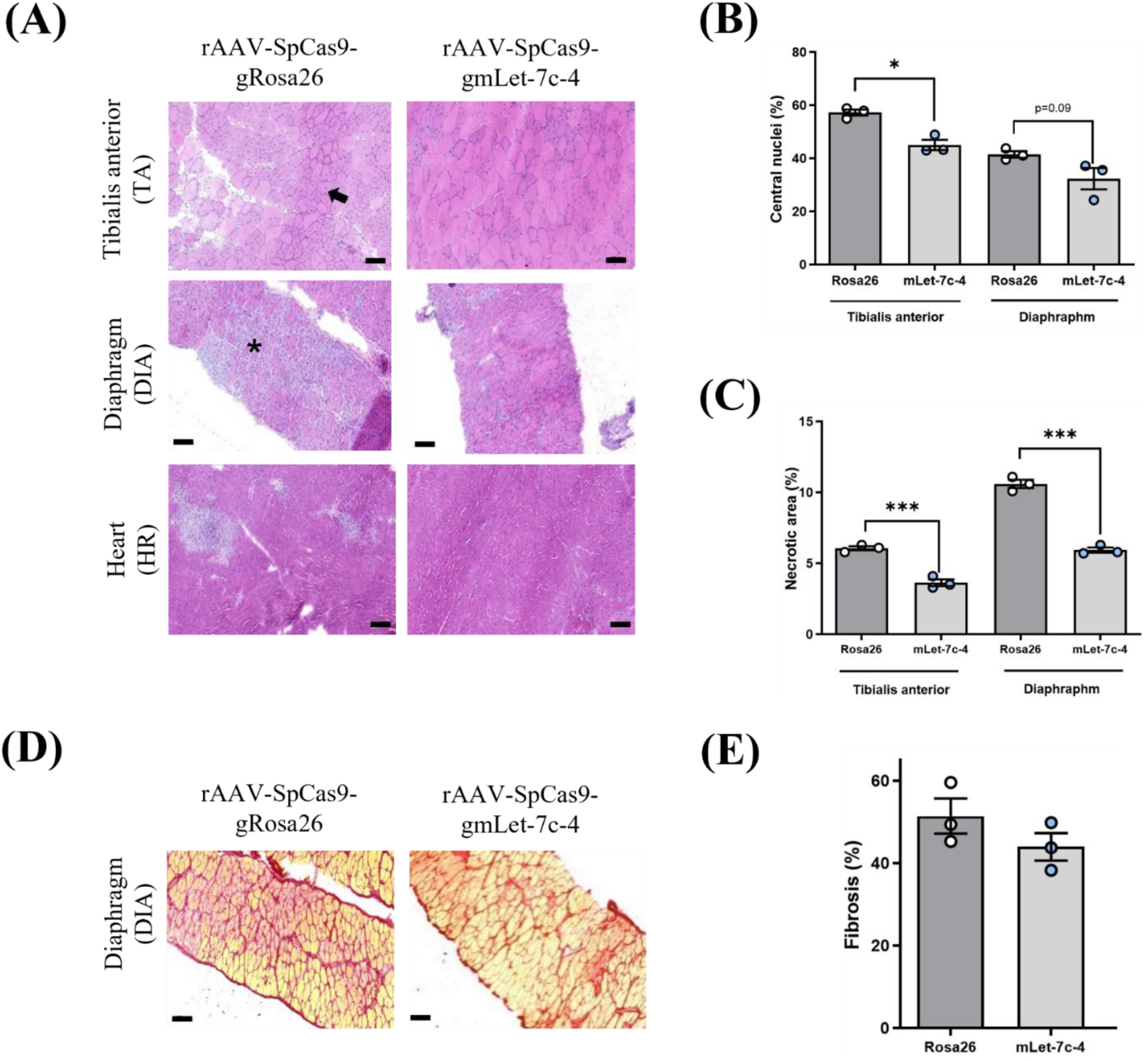
Cas9/gRNA AAV treatment improves histological defects in mdx mice. **(A)** Representative images of hematoxylin and eosin stained transverse muscle sections of TA, DIA and HR (9 weeks of age) in rAAV-gRNA mLet-7c-4 and rAAV-gRNA Rosa26 (control) treated mdx mice showing necrotic areas (black stars) and regenerating fibers (black arrows). Scale bar: 100 µm. **(B)** TA and DIA muscle from mice treated with rAAV-gmLet-7c-4 showed a decrease in centrally nucleated fibers compared to control/rAAV-gRosa26. Bars are mean ± SEM of n = 3 per condition. Dots represent each mouse. p < 0.05. **(C)** The necrotic muscle area in TA, DIA and HR of mice treated with rAAV-gmLet-7c-4 is decreased compared to the control/rAAV-gRosa26 group. Bars are mean ± SEM of n = 3 per condition. Dots represent each mouse. ***p < 0.001. **(D)** Representative images of sirius red staining of diaphragm in rAAV-gmLet-7c-4 and rAAV-gRosa26 (control) treated mdx mice. **(E)** Sirius red quantification indicated that rAAV-gmLet-7c-4 treatment reduced the amount of collagen infiltration (fibrosis) by 10% in diaphragm muscle. Bars are mean ± SEM, n = 3 per condition. Dots represent each mouse TA, tibialis anterior; DIA, diaphragm; HR, heart.

In summary, our data demonstrate the in vivo potential of our CRISPR-Cas9 strategy to upregulate utrophin expression and consequently to improve dystrophic defects in skeletal, respiratory and cardiac mdx muscles.

## DISCUSSION

Utrophin upregulation is an attractive therapeutic avenue, applicable to all DMD patients, independently of their genetic defect, able to compensate for the lack of dystrophin in DMD (*18*). In the present manuscript, we describe the in vitro and in vivo potential of an innovative gene editing strategy to upregulate the expression of the full-length endogenous utrophin in human and murine muscle cells as well as in the mdx mouse model. Several miRs, such as miR-133b, miR-150, miR-196b, miR-296-5p and Let-7c, were previously described to have their BSs clustered in a 500bp sequence within the utrophin 3’UTR and be able to post-transcriptionally repress utrophin expression (*42*). We first confirmed, using a Cas9/gRNA gene editing strategy and reporter plasmids, that miR Let-7c BS is the major inhibitory site to disrupt to upregulate utrophin mRNA and protein levels by 3-fold in human DMD myoblasts. Nevertheless, current conventional 2D cultures fail to recapitulate key aspects of the disease observed in humans (*75*). To address this major limitation we investigated the benefits of our approach in 3D engineered muscles generated from immortalized human DMD myoblasts (*65, 69*). After Cas9/gRNA treatment, we observed a significant increase of utrophin levels associated with an amelioration of calcium dysregulation as well as a higher number of DMD 3D muscles responding to electrical stimulation with measurable contractions. Considering these promising data, together with the lack of major gRNA associated off-targets, we decided to investigate the potential of this gene editing strategy in vivo in mdx mice. After choosing an efficient gRNA targeting the murine miR Let-7c BS, we designed a two rAAV strategy to deliver the Cas9/gRNA components in vivo and opted for a systemic injection administration route, which is currently explored in DMD clinical trials. In treated mdx mice, we observed good AAV transduction and sufficient editing to obtain a 2-fold utrophin upregulation resulting in significant histological benefits on muscle regeneration, necrosis/inflammation and fibrosis defects.

Anti-sense approaches such as 2’AON and P-PMO masking the BS of miR Let-7c were previously described to raise utrophin levels and improve biochemical and morphological DMD manifestations in mice; however, improvements were limited and the strategy required weekly administration (*43, 55*). CRISPR-Cas9, instead, can induce permanent DNA alteration with a single administration, providing a crucial advantage over transient anti-sense approaches. Recently, two different groups described the use of a dual gRNAs strategy to delete the utrophin 3’UTR sequence containing the miR BS cluster and reported a 1.5-2-fold utrophin upregulation in DMD immortalized myoblasts and DMD human induced pluripotent stem cells (*55, 56*). Using reporter plasmids, we observed that deletion of miR Let-7c alone results in even higher utrophin upregulation, suggesting that miR Let-7c is the main driver of utrophin repression on the 3’UTR and/or that some sequences in the miR cluster region contain additional elements important for mRNA stability or translation. Intriguingly, we also observed that indels generated with a single gRNA targeting miR hLet-7c BS were as efficient as the full BS deletion in upregulating utrophin, indicating that even a single nucleotide change can have a strong impact on miR binding. The differences observed using multiple gRNA targeting Let-7c BS reflect the importance of assessing both gRNA activity, i.e indel efficiency, as well as editing outcome, i.e. indel pattern, with indels in the miR seed regions being more disruptive (*76, 77*). Overall, our single gRNA approach seems more efficient than the described dual gRNA strategies, which simplifies delivery and potentially induces less DNA damage and off-targets.

For mouse studies, we performed systemic injection of two AAV9 carrying Cas9 and gRNA. Our indels rate, ranging between 5% (DIA), 12% (TA) and 21% (HR), were sufficient to upregulate utrophin and ameliorate DMD histopathological phenotype; however, we envision that new AAV serotypes, such as MyoAAV (*78*), reported to achieve better transduction of skeletal and cardiac muscles, will allow higher indels, utrophin upregulation and therapeutic benefits.

However, several aspects need to be improved before this approach can be translated to the clinics. Firstly, the large 4.1 kb size of SpCas9 endonuclease dictates the use of a dual rAAV strategy, which requires higher vector doses and potentially results in more side effects linked to rAAV injection and stronger immune response against the rAAV capsid (*79*). An all-in-one rAAV encoding for single gRNA, regulatory elements and smaller Cas9 orthologs such as SaCas9 (3.2kb) (*80*), CjCas9 (2.9kb) (*81*) or Nme2Cas9 (3.2kb) (*82*) may be required. Although we propose to use a single gRNA, which has less risk compared to dual gRNA approaches, off-targets generated by double-strand DNA breaks (DSB) introduced during Cas9/gRNA editing are a major concern. In addition, on-target DSB may lead to large deletions, complex rearrangements, chromothripsis or aneuploidy and chromosomal truncations (*83–85*). Hence more controlled novel and safer CRISPR-Cas9 based tools would be desirable in the future for translation (*63*). Since we demonstrated that a minimal modification of the mRNA Let-7c BS, e.g. a +1 nucleotide insertion, is sufficient to block miR Let-7c mediated utrophin repression, we expect our strategy to be easily adapted for base and prime editing tools, which do not induce DSBs (*86–88*). In addition, these systems could be used to simultaneously disrupt multiple utrophin repressor BS. As an example, the homeobox protein engrailed-1 (EN1) (*89*) and the Ets-2 (*90*) factors were both described to repress utrophin promoter activity. Recently, the binding of Poly (C) binding protein 2 (PCBP2) to the 5’UTR of utrophin was described to downregulate utrophin-A expression (*91*). Therefore, disruption of the BS of these transcriptional repressors on the utrophin promoter and 5’UTR, alongside the Let-7c BS on the utrophin 3’UTR, may result in additive or synergic utrophin upregulation. Recently, dystrophin and utrophin were described to co-localize, to a certain extent, at the same muscle membrane (*38*). Therefore, editing strategies that restore dystrophin and upregulate utrophin could also be combined. In summary, we provide in vitro and in vivo evidence of a novel gene editing strategy to upregulate utrophin expression via disruption of miR Let-7c BS, amenable for all DMD mutations.

The described therapeutic potential of editing RNA regulatory elements for regulating gene expression combined with high-throughput techniques for their identification (*92–94*) will pave the way for modulating others disease modifier genes associated with DMD, such as follistatin (*95*) or Jagged-1 (*96*), as well as other diseases (*97–99*).

## MATERIALS AND METHODS

### Ethical statement

All animal procedures were performed in accordance with the European guidelines for the human care and use of experimental animals, and animal experimentations were approved by the Ethical Committee for Animal Experimentation C2AE-51 of Evry under number APAFIS#29497-2020102611378971 v2 and DAP 2020-001-B. All C57BL/10ScSn-Dmdmdx/J (BL10/mdx) male mice were bred in CERFE (Experimental Functional Research Exploration Center) facility, Génopole. Work with human cells in the Dr Tedesco laboratory was performed under approval of the NHS Health Research Authority Research Ethics Committee reference no. 13/LO/1826; IRAS project ID no. 141100.

### Cell cultures

Human DMD myoblasts were maintained in Smooth Muscle Cell Growth Medium (C-23060, PromoCell) supplemented with 20% fetal bovine serum gold (PAA) and 1% penicillin– streptomycin (Invitrogen). Wild-type murine C2C12 were maintained in DMEM (Invitrogen) supplemented with 20% fetal bovine serum gold (PAA) and 1% penicillin–streptomycin (Invitrogen). Cells were maintained at 5% CO2 at 37°C. At 70% of myoblasts confluence, myotubes differentiation was induced using DMEM + 2% horse serum. HEK293 cells were cultured in Advanced DMEM (Life Technologies) supplemented with 10% FBS, 2 mM GlutaMax (Life Technologies), and penicillin/streptomycin at 37 °C with 5% CO2.

### gRNAs design

The guides targeting the miR BS were chosen based on their proximity to the mutation intended for editing and designed on the basis of the most active sgRNAs as computationally predicted by the online Benchling Tool (*100*). All sgRNAs with a predicted activity score greater than 0.30 were next analyzed by the CRISPR Design tool and ranked according to the least possible number of potential off-target sites (*101*). All gRNA sequence used are specified in the Supplemental data Table S1.

### Nucleofection

Chemically modified single guide RNA (Synthego) were diluted following manufacturer’s instruction. Ribonucleoprotein complexes were formed with sgRNA and 30 pmol of Streptococcus pyogenes Cas9 protein (gift from Dr J.P Concordet) (ratio 1:2) (*102*). 2.5 x 10^5^ hDMD or C2C12 cells per condition were transfected with RNP using P5 Primary Cell 4D-Nucleofector X Kit (C2C12 program). Culture medium was replaced the following day, and cells were harvested for DNA, RNA or protein analysis 7 days after electroporation.

### DNA analysis

Genomic DNA was extracted with the QuickExtract™ DNA Extraction Solution (Lucigen, Middelton, WI, USA). 50 ng of genomic DNA were used to amplify the region that spans the cutting site of each gRNA using KAPA2G Fast ReadyMix (Kapa Biosystem, Wilmington, MA, USA). After Sanger sequencing (Genewiz, Takeley, UK), the percentage of insertions and deletions (InDels) was calculated using TIDE (*103*) or ICE software (*104*).

### RNA Extraction and RT-qPCR

Total RNA was purified using RNeasy Micro kit (Qiagen, Hilden, Germany). RNA was reverse-transcribed using Transcriptor First Strand cDNA Synthesis Kit (Roche, Basel, Switzerland). qPCR was performed using Maxima Syber Green/Rox (Life Scientific, Thermo-Fisher Scientific, Waltham, MA, US). Utrophin A (forward primer 5’-ACGAATTCAGTGACATCATTAAGTCC-3’, reverse primer 5’ ATCCATTTGGTAAAGGTTTTCTTCTG-3’) mRNA expression levels were normalized using human GAPDH as a reference gene (NM_002046.6) and represented as fold changes (2∧ ΔCt) relative to the control. PCR amplification efficiency was determined by applying linear regression analysis to the exponential phase of the amplification curve of each PCR reaction using the LinRegPCR software. No reverse transcriptase (non-RT) and no template control (NTC) reactions were used as negative controls in each 40-cycle PCR run (Cq values NTC = undetermined, non-RT = undetermined).

### Protein analysis

Human and mouse cells and muscles were homogenized on ice in RIPA buffer (R0278-50ml, Sigma-Aldrich) supplemented with protease inhibitors (P8340, Sigma-Aldrich). Following BCA quantification, 10μg of total protein were heat-denatured for 5 minutes at 100°C before loading onto NuPAGE 3– 8% TRIS Acetate Midi Gel (Novex, Life Technologies) and transferred to PVDF membranes (Millipore). Membranes were blocked for 1 hour with Odyssey Blocking buffer (926-41090; LI-COR; USA) and then incubated with the following primary antibodies overnight at 4°C: mouse anti-utrophin (1:100, Utrophin (84A), SC-33700, Santa Cruz Biotechnology), rabbit anti-gapdh (1:5000, MAB374, Sigma Aldrich), mouse anti-α-actinin (H-2) (1:1000, sc-17829, Santa Cruz Biotechnology), rabbit anti-vinculin (1:2000, ab73412, Abcam). The Odyssey Imaging System (LI-COR Biosciences; USA) was used to read infrared fluorescence of the secondary antibodies and the Image Studio Lite software (LI-COR Biosciences; USA) to quantify target proteins relative to gapdh, α-actinin or vinculin.

### Reporter plasmid contruction and transfection

The utrophin 3’UTR is based on the human utrophin UTRN-001 (ENST00000367545.7) sequence. All utrophin 3’UTR reporter constructs were generated by GenScript Biotech (Leiden, Netherlands) and inserted in the pEZX-GA02 Gaussia luciferase (Gluc) and secreted alkaline phosphatase (SEAP) reporter cloning vector (ZX-104, Genecopoeia) downstream of the Gaussia luciferase reporter gene. Plasmids were controlled by enzymatic digestion and Sanger sequencing. Following transformation in XL-10 bacteria, plasmid preparation was performed using NucleoSpin Plasmid kit (740588.50, Macherey Nagel) and following manufacturer recommendations. To study the impact of utrophin 3’UTR variants on the Gaussia Luciferase reporter gene expression, hDMD Δ52 myoblast were seeded in 96-well plates at 10,000 cells/well. The day after, cells were transfected using Lipofectamine™ 3000 (L3000008, ThermoFisher) as transfection agent. Briefly, 100ng of the pEZX-GA02-3’UTR variant and the 0.2ul of P3000 reagent were diluted in 5ul of Optimen prior to be gently mixed with 0.3ul of Lipofectamine 3000 diluted in 5ul of Optimen. After 15 min of incubation at room temperature, the mixture was diluted with serum-free culture medium to a final volume of 100µl. Experiments were done in triplicate. Forty-eight hours after transfection, supernatant was collected for enzymatic dosages.

### Enzymatic dosages

Gaussia luciferase activity was measured using the following protocol: cell culture medium was collected and diluted in PBS1X using a 1:10 dilution. 50ul of diluted supernatant were distributed in white 96-well OptiPlate. 11ul of Coelenterazine (C3230-50UG, Sigma Aldrich) were diluted in 5.5ml of PBS1X and automatically distributed. Luciferase light units were measured using the EnSpire Multimode Plate Reader (Perkin Elmer, Courtaboeuf, France). The transfection efficiency was controlled by quantification of the SEAP using the Phospha-Light™ SEAP Reporter Gene Assay System (T1015, ThermoFisher) and a 1:20 dilution of the supernatant. Gaussia luciferase value were normalized by SEAP measurements. All conditions were tested in triplicate.

### Guide-Seq

DNA libraries preparation for GUIDE-seq analysis was performed as previously described (*105, 106*). The optimal dsODN concentration based on integration efficiency by DECODR v3.0 analysis (https://decodr.org/) and cell viability by cell counting were determined after nucleofection of 2.105 HEK293T cells with different dsODN concentrations (60, 80, 100 and 120 µM). 80 µM of dsODN results in a 10% integration and was used for further transfections together with gRNA Let-7c-1 and Cas9 RNP at a 2:1 molar ratio. After 4 days in culture, DNA was isolated with QIAamp DNA mini kit using standard protocols (QIAGEN). DNA fragments of 400–900 bp were generated by sonication and subsequently ligated to adaptors, followed by two steps of DNA amplification by utilizing KAPA HyperPrep Kit (Roche KK8504). Then, the libraries were purified and measured by qubit, the concentrations evaluated by qPCR KAPA libraries quantification kit (Roche) and the average bp length was estimated by Tape Station bianalyzer 2100 (Agilent). Finally, the libraries were pooled, diluted to reach 4nM and loaded into MiSeq flow cell. Demultiplexing, PCR duplicate consolidation, cleavage on target site recognition Let-7 ATGGATCTGAGGTAGAAAGGTGG:“chr6:144852595”, off-target activity identification, and visualization was performed with the GUIDE-Seq Analysis pipeline from Bushman lab https://github.com/cnobles/iGUIDE using the hg38 human genome as reference.

### Amplicon-Seq

Amplicons were generated on genomic DNA extracted from control and RNP treated 293T and hDMD, to quantify on- and off-target indels generation, or on different tissues (HA, DIA, TA) of mice treated with AAV gRNA mLet7c-4, to quantify on-target indes. Amplicons were prepared by two round amplifications of genomic DNA. The first amplification was specific for *each target*: a 50-μl PCR mixture containing 27 μl nuclease free Water, 10 μl of PCR GC buffer, 2.5 μl forward primer (10 μM), 2.5 μl reverse primer (10 μM), 1.5 μl DMSO, 1 µl of dNTPs (10 mM), and 200 ng of template DNA was generated. The PCR program was as follows: (i) 3 min at 98°C, (ii) 32 cycles of 10 s at 98°C, 10 s at the specific primer Tm, and 15 s at 72°C, and (iii) 5 min at 72°C. The amplicons were purified using an Agencourt AMPure XP beads kit (Beckman Coulter, Brea, CA, USA) and used in the second round of amplification with specific barcoded primers. The final amplicons are gel purified using NucleoSpin Gel and PCR clean up from Macherey Nagel, following the manufacturer’s instructions. The final products were eluted in Tris (10 mM, pH 8.5) buffer. After purification, the PCR products were pooled in equimolar concentrations and were delivered to the NGS facility core (Institut Imagine Paris, France) for 100 paired ends sequencing on an Illumina NovaSeq instrument. The Amplicons were cleaned, quality filtered, demultiplexed and processed using CRISPResso V2 online tool (https://crispresso.pinellolab.partners.org/submission). The sequencing data were deposited (https://dataview.ncbi.nlm.nih.gov/object/PRJNA941216?reviewer=ci5vg8qgorak8qnodj1mr17ln0).

### Generation of 3D artificial muscle constructs

3D artificial muscles were generated as previously reported (*68, 69*) using 10% Matrigel (BD, 356230) and 3.5 mg/ml human fibrinogen (Tissucol Duo 500, Baxter; Sigma-Aldrich, F8630) polymerized by adding 3 U/ml thrombin (Biopur, 10-13-1104; Sigma-Aldrich, T7326) to the solution. 1 x 106 immortalized myoblasts were combined with the aforementioned matrix cocktail to form a 3D artificial muscle hydrogel (of total volume 120μl) which were polymerized between two silicone posts (EHT Technologies, GmbH Hamburg). After polymerization the 3D artificial muscle gels were cultured at 37°C with 5% CO2 supplementing the culture medium (PromoCell, C-23060) with 33μg/μl aprotinin (Sigma, A3428) to prevent fibrinogen degradation. To induce myogenic differentiation, 48 hours after polymerization in culture medium, the 3D artificial muscle gels were transferred into differentiation medium (PromoCell, C-23061) supplemented with aprotinin. The gels were maintained in differentiation media with media change every alternate day until day 14 from gel polymerization.

### Immunostaining of whole 3D artificial muscles

For immunostaining, 3D artificial muscles were fixed with buffered 1% paraformaldehyde (PFA) (ThermoScientific, JI9943.K2) overnight at 4°C followed by 6 hours of blocking at 4°C [10% FBS (Gibco, 10270-106), 1% BSA (Sigma, A7906-100G), and 0.5% Triton X-100 (Sigma, T8787-250ml) in 0.05 M Tris-buffered saline (TBS)] before immunolabeling with mouse anti-Utrophin (8A4) (SantaCruz, SC 3370), mouse anti-myosin heavy chain (DSHB, MF20, RRID: AB_2147781) and mouse anti-titin (DSHB, 9 D10, RRID: AB_528491) primary antibodies overnight at 4°C in TBS, 1% BSA, (Sigma, A7906-100G) and 0.5% Triton X-100 (Sigma, T8787-250ml). The next day, the 3D artificial muscles were washed six times with TBS, with each wash for one hour at RT and then incubated overnight with Hoechst 33342 (Sigma, B2261) and species-specific secondary antibodies Alexa Fluor 488, 546, and 647 (Thermo Scientific). The following day, 3D artificial muscles were again washed six times with TBS and embedded in mounting medium (Dako, S3023A) on glass slides for downstream microscopic analysis. A confocal microscope (Leica bio systems) was used for imaging.

### Processing and Immunostaining of 3D artificial muscles Transverse section

Prior to cryo-embedding of 3D artificial muscles for transverse section, the muscles were fixed with buffered 1% paraformaldehyde (PFA) (ThermoScientific, JI9943.K2) overnight at 4°C. Thereafter, the muscles were treated with sucrose (Sigma, S0389) solution gradient for specific time durations: 10% sucrose -16hr, 15% sucrose -16hr and 30% sucrose -16hr. Post sucrose treatment, the muscles were embedded with Tissue-Tek OCT solution (Sakura Finetek, 4583) on dry-ice. The embedded 3D artificial muscles were transverse sectioned onto glass slides using a cryostat (Leica). The muscle sections were immuno-stained using the mentioned procedure. The slides with muscle sections were washed twice with PBS (Sigma, P4417) for 3 mins at room temperature (RT). Then the sections were permeabilized with PBST (PBS with 0.5% Triton X (Sigma, T8787-250ml)) for 15 mins at RT and followed by blocking with 10% BSA in PBST at RT for 1hr. Subsequently, the muscle sections were incubated overnight with primary antibody diluted in 5% BSA (Sigma, A7906-100G) at 4°C. After primary antibody incubation, the sections were washed four times with PBST, with each wash for 3 mins at RT. Post PBST wash, the sections were incubated with Hoechst 33342 (Sigma, B2261) and species-specific secondary antibodies (Thermo Scientific) at RT for 2 hrs. Thereafter, the muscle sections were washed four times with PBST, with each wash for 3 mins at RT and thereafter mounted for downstream microscopy.

### Calcium transient assay for 3D artificial muscles

Artificial muscles were mounted on PDMS posts placed into a 24 well plate and incubated with 2 ml Opti-MEM basal media without phenol red containing Fluo-4AM (Thermo scientific, F14201) as per manufacturer’s instructions for 30mins. Thereafter muscles were washed with Opti-MEM basal media without phenol red for 10mins and processed for calcium transient upon assay electrical stimulation. EHT carbon electrodes (EHT Technologies, Germany) were carefully placed on either side of the muscle without touching the muscle. Electrodes were temporarily attached to the 24 well plate using Blu Tack®. The entire set was placed on Zeiss 880 confocal LSM stage for fluorescence excitation at 488 nm and emission collected at >530 nm and prepare to acquire as a time series at 33 fps. Fields with aligned myotubes were selected and imaged using a 10x objective. The electrical stimulator connected to the electrodes was adjusted to 0.5Hz and voltage setting of 5V and 10V as required. Before stimulation, 10sec of baseline recording was done without electrical stimulation, followed by 30sec of electrical stimulation at 5V, followed again by 30sec 10V. Export time series from microscope and open the time series file using FiJi software for downstream analysis, wherein z-axis profile is plotted for the mean intensity signal.

### 3D artificial muscle contractility assay

Artificial muscles mounted on PDMS posts were placed into a 24 well plate, each with 2 ml muscle differentiation media. The 24 plate along was then placed onto an elevated plate holder equipped with a Dino-Lite camera (Dino-Lite digital microscopy, The Netherlands) positioned underneath. Carbon electrodes (EHT Technologies, Germany) were carefully placed on either side of the muscle without touching the muscle, temporarily attached to the 24 well plate using Blu Tack®. The electrical stimulator connected to the electrodes was set at 0.5Hz and 5-10V as required. The recorded contraction activity (.avi) was analysed using MUSCLEMOTION (*107*) plugin on ImageJ. Before stimulation, 10sec of recording was done without electrical stimulation to measure baseline activity.

### AAV cloning, production and quantification

The AAV encoding for SpCas9 under the control of the CMV promoter (pX551-CMV-SpCas9) was a gift from Alex Hewitt (Addgene plasmid # 107024). For the AAV encoding for sgRNA and GFP, we started from the pAAV-U6-sgRNA-CMV-GFP, which was a gift from Hetian Lei (Addgene plasmid # 85451). First, we replaced the sgRNA scaffold to insert the optimized one (*108*) and then we cloned the gRNA protospacers of interest by Sap I digestion (*109*). Each plasmid was checked by digestion and Sanger sequencing. All reagents and detailed sequences information are available upon request. AAV vectors were produced following a triple transfection protocol using 293 cells in suspension and purified by affinity chromatography. To determine vector copy number, genomic DNA was isolated from TA, DIA and HR muscles using the KingFisher Flex device and the NucleoMag Pathogen kit (744210.4, Macherey-Nagel). Briefly, AAV genome copy number in tissue samples was quantified using real-time TaqMan PCR analysis (ABI 7500). The primers and probe set were directed against the CMV promoter of the rAAV9-SpCas9 vector (CMVFor: 5’ catcaatgggcgtggatagc 3; CMVRev: 5’ ggagttgttacgacattttgg 3’). PCR conditions were 60°C for 2 min, 95°C for 10 min, and 40 cycles of 95°C for 15 sec and 60°C for 1 min using 100 ng of genomic DNA in 1× TaqMan Universal PCR master mix (Life Technologies, Carlsbad, CA) in a 10 μl volume. A plasmid containing the CMV promoter was used to generate a standard curve at a range from 1× 1010to × 108.

### Mouse treatment

Four-week-old mdx mice were administered by tail-vein injection with a total 10^13^ vector genomes of rAAV9-CMV-Cas9 and rAAV9-gRNA, with a 1 to 5 ratio. For histological and molecular analysis of mouse tissues, specimens were collected immediately after animals were sacrificed by cervical dislocation, snap frozen in liquid-nitrogen-cooled isopentane and stored at −80°C.

### Histological analyses

Tibialis anterior (TA) muscle transverse cryosections (8 µm thickness) were prepared from frozen muscles, air dried, and stored at −80°C. Mouse sections were processed for Hematoxylin-Eosin staining as previously described (*39*). Whole muscle sections were visualized on an Axioscan Z1 automated slide scanner (Zeiss, Germany), using the ZEN2.6 SlideScan software and a Plan APO 10×0.45 NA objective. The proportion of centrally nucleated fibres was determined by analysing the H&E images of the whole muscle section. Areas of necrosis were quantified based on the DMD_M.1.2.007 MDC1A_M.1.2.004 TREAT-NMD SOPS and performed with the Fiji ImageJ 1.49i software on the TA sections. Mouse sections were processed for Sirius Red staining as previously described (*40*). Fibrose quantification was processed in QuPath (*110*) using the pixel classifier feature. A machine learning model based on random forest method was trained using a ground truth data corresponding to selected regions containing fibrosis defined as “Fibrosis” class, regions containing muscles defined as “Muscle” class and regions containing no tissue or fat defined as “Blank” class. For each slice, a region containing the muscle slice was defined using the qupath brush and wand tools excluding visually parts containing artifacts like tissue folding. The pixel classifier model was applied to the region and regions of each class were predicted. and the total areas of each class were extracted. Ratio of fibrosis was calculated by dividing the total “Fibrosis” area with the total area of the muscle slice.

### Immunofluoresence

Frozen transverse muscle sections were fixed 10 min in acetone, then blocked in M.O.M.® (Mouse on Mouse) (BMK-2202, Vector Laboratories) for 30 min and incubated with the mouse monoclonal anti-utrophin (1:50, SC-33700) and rat anti-laminin-α2 (1:50, SC-59854) primary antibodies overnight at 4°C. Sections were next washed in PBS and incubated with suitable Alexa Fluor secondary antibodies for 1 h at room temperature. Whole muscle sections were visualized on an Axioscan Z1 automated slide scanner (Zeiss, Germany), using the ZEN2.6 SlideScan software and a Plan APO 10×0.45 NA objective. To quantify utrophin signal, fiber cytoplasmic region enclosed within the membrane laminin-α2 staining are segmented by morphological segmentation after contrast enhancement and artefact filtering (FiJi software 2.0.0-rc/1.52p, Morpholib plugin v 1.4.1). Fibers are enlarged according to the magnification to capture the membrane region. The fluorescence intensity in each object (fibers, fibers membrane) is measured for each channel together with fibers shape and size. Positive fibers for any channel are detected based on the fluorescence distribution of negative control slices or slices from known negative condition.

### Statistical analysis

Results were analyzed using Prism (GraphPad Software, Inc.). For statistical significance, we performed the Student’s t test with a two tailed distribution assuming equal or unequal sample variance depending of the equality of the variance (F-test) and Bonferroni correction. Data are presented as mean ± SEM (standard error of mean), with n indicating the number of independent biological replicates used in each group for comparison. Differences were considered significant at (*) p<0.05; (**) p<0.01 and (***) p<0.001.

## List of Supplementary Materials

Fig. S1. Indel profiles of gRNAs disrupting miR BS

Fig. S2. Myotube differentiation of edited hDMD Δ52 myoblasts

Fig. S3. Characterisation of 3D hDMD artificial muscles

Fig. S4. hLet-7c-1 gRNA off target analysis

Fig. S5. Indel profiles after rAAV-mLet-7c-4 treatment in mdx mice

Table S1. gRNA sequences used in this study

## Acknowledgments

The authors are Genopole members, first french biocluster dedicated to genetics, biotechnologies and biotherapies. We acknowledge the Genethon “Vector Core Facility” for the production of the rAAV, Genethon “Functional Evaluation Facility” for help with mice experimentation and histology/immunofluorescence and Genethon Bioinformatics Platform for image analysis. We are grateful to Genethon “Imaging and Cytometry Core Facility” for technical support, to Ile-de-France Region, to Conseil Départemental de l’Essonne (ASTRE), INSERM and GIP Genopole, Evry for the purchase of the equipments. and the Imagine “Genomic Platform” for Guide-seq and amplicon sequencing. We also acknowlege the CERFE for the animal work. We acknowledge Pr. Vincent Mouly for the hDMD Δ52 myoblasts cell line, Dr JP Concordet for Cas9 protein production (supported by ANR-II-INSB-0014). We thank the members of the “Therapeutic genome editing” group for fruitful discussions and experimental help. We gratefully acknowledge the Conseil Général de l’Essonne (ASTRES) and Genopole Research in Evry for financial help for the purchase of equipment. SD thanks Valentina Lionello for help and assistance with artifical muscle functional tests.

## Funding

Agence Nationale de la Recherche grant HemoLEn ANR-20-CE17-0016-0 (MA)

Agence Nationale de la Recherche grant PEMGeT ANR-22-CE17-0028-02 (MA)

Génopole (SG)

AFM-téléthon (MA)

INSERM (MA)

University of Évry Val d’Essonne (MA)

Paris Ile de France Region – “DIM Thérapie génique” initiative (MR)

European Research Council grant 759108 – HISTOID (FST)

National Institute of Health Research UK grant CL-2018-18-008; *the views expressed are those of the authors and not necessarily those of the National Health Service, the NIHR, or the Department of Health* (FST)

The Francis Crick Institute

Cancer Research UK

UK Medical Research Council The Wellcome Trust (FC001002)

## Author contributions

Conceptualization: SG, MA

Methodology: SG, SD, FM, FA, MR

Investigation: SG, SD, FM, FA, MR, MA

Supervision: SG, MA

Writing—original draft: SG, MA

Writing—review & editing: SD, FM, FA, MR, AC, IR, GR, FST

## Competing interests

S.G and M.A are inventors on a patent application related to this work filed by the Genethon (PCT/EP2021/076882; 10243723, filed 29 September 2021). FST has received speaker and consultancy honoraria from Takeda, Sanofi Genzyme and Aleph Farms (via UCL Consultants). All other authors declare that they have no competing interests.

## Data and materials availability

The authors declare that all data supporting the findings of this study are available in the main text or the supplementary materials. The Guide-seq and Amplicon-seq data that support the findings of this study have been deposited (https://dataview.ncbi.nlm.nih.gov/object/PRJNA941216?reviewer=ci5vg8qgorak8qnodj1mr17ln0). Any other relevant data is available upon reasonable request.

## Notes

### Competing Interest Statement

S.G and M.A are inventors on a patent application related to this work filed by the Genethon (PCT/EP2021/076882). FST has received speaker and consultancy honoraria from Takeda, Sanofi Genzyme and Aleph Farms (via UCL Consultants). All other authors declare that they have no competing interests.

